# On-lamella super-resolution cryo-CLEM for cryo-ET enabled by vacuum-free ultra-stable cryogenic fluorescence microscopy

**DOI:** 10.64898/2026.04.14.717675

**Authors:** Julian Falckenhayn, Vi Quint Duong, Neeraj Prabhakar, Iain Harley, Enoch Lok Him Yuen, Tolga O. Bozkurt, Stephen D. Carter, Vojtěch Pražák, Rainer Kaufmann

## Abstract

Cryogenic correlative light and electron microscopy (cryo-CLEM) combines specific fluorescence labelling of proteins inside cells with structural information at the angstrom-level. The introduction of super-resolution fluorescence methods in the field of cryogenic fluorescence microscopy is a necessary step to bridge the large resolution gap between the different imaging modalities. However, there are many challenges hindering the full potential of cryogenic super-resolution correlative light and electron microscopy and seamless integration with structural cell biology. One of the main limiting factors is a lack of dedicated cryogenic fluorescence microscopy systems with sufficient mechanical stability to enable the collection of high-quality super-resolution data and full compatibility with vitrified specimens for cryo-electron tomography. Here, we address this by developing a vacuum-free ultra-stable cryogenic optical microscope (VULCROM). VULCROM is a dedicated super-resolution cryo-CLEM (cryo-SR-CLEM) setup that combines the stability of a vacuum-insulated cryostat with the flexibility and modularity of an open microscopy system. We demonstrate that VULCROM enables detailed investigations of single-molecule cryo-photo-physics across timescales spanning milliseconds to hours. We furthermore demonstrate its suitability for routine cryo-SR-CLEM with a resolution in the 10 nm range in distinct vitrified biological specimen types. We resolve the nanoscale architecture of YFP-labelled PML bodies within the nucleus of mammalian cells and the distribution of ATG9-eGFP in its cellular structural context in a cryo-lift-out lamella of *N. benthamiana* plant tissue. Owing to its vacuum-free design, VULCROM can be readily adapted for diverse correlative workflows and other cryo-light microscopy applications.

## Introduction

Cryogenic super-resolution fluorescence microscopy (cryo-SR-FM) bridges the resolution gap between conventional cryo-FM and cryogenic electron microscopy (cryo-EM) ^1, 2, 3, 4, 5^. Cryo-SR-FM is an important development for correlative cryo light and electron microscopy (cryo-CLEM) – a workflow in which the highly specific information of fluorescence labelling is used to locate and validate the identity of protein complexes in vitrified specimens in the context of the structural information provided by cryo-EM ^1, 2, 3, 4^. This is of particular interest for cryogenic electron tomography (cryo-ET) which, combined with sub-tomogram averaging, can resolve the structure of protein complexes down to the angstrom-range even in the crowded environment of a cell ^6, 7, 8^. The principal feasibility of cryo-SR-FM ^9, 10, 11, 12, 13^ and the combination with cryo-ET has been demonstrated in a variety of approaches ^9, 14, 15, 16, 17^. However, performing cryo-SR-FM still poses challenges that have not yet fully been overcome. The major remaining challenges include (i) reaching sufficiently high mechanical stability of the cryo-microscopy setup; (ii) limited photon detection efficiency due to the restricted numerical aperture (NA) of objective lenses suitable for cryogenic optical imaging; (iii) incomplete understanding of single-molecule photo-physics under cryo-conditions and non-ideal photo-switching/blinking behaviour; and (iv) devitrification of the specimen caused by the high laser intensities required for photo-switching/blinking ^3, 4, 5^. A dedicated cryo-SR-FM setup must address two further practical requirements for successful cryo-ET data collection. First, a sample transfer mechanism is needed that maintains the vitrified state of the specimen throughout. Second, build-up of ice contamination from atmospheric humidity, whether during sample transfer or at any point during imaging, must be minimised.

Under cryo-conditions, single fluorescent molecules can exhibit extended on-states lasting seconds to minutes, allowing the accumulation of sufficient photons for single-molecule localisation precision in the sub-10-nm range ^11, 17^, and for a limited number of molecules per diffraction volume even angstrom-range ^13, 18, 19^. However, the slow photo-switching/blinking kinetics at low temperature mean that cryogenic single-molecule localisation microscopy (cryo-SMLM) datasets can take hours to days to acquire ^11, 17^, placing stringent demands on the long-term mechanical and thermal stability of the microscopy setup. Early cryo-SMLM approaches based on cryo-chambers attachable to standard microscope stages enabled first proof-of-concept demonstrations; however, limiting achievable resolution due to remaining instabilities ^9, 10, 16, 17^. Vacuum-insulated cryostats have since delivered the highest stability ^11, 13, 18, 20^, but at the cost of flexibility, modularity and increased complexity ^1, 4^.

Here, we introduce the vacuum-free ultra-stable cryogenic optical microscope (VULCROM). VULCROM is a dedicated cryo-SR-FM system that combines the flexibility and simplicity of a vacuum-free system with the thermal and mechanical stability of a vacuum-based cryostat. VULCROM is fully compatible with standard cryo-ET workflows, enabling cryo-SR-CLEM for applications in structural cell biology, while providing sufficient stability for single-molecule cryo-photo-physics investigations across millisecond to hour timescales. We first demonstrate the general performance of the VUCLROM setup by characterising its stability and optical capabilities. We then demonstrate the capacity of VULCROM to capture the full range of single-molecule photo-switching behaviour under cryo-conditions, from fast millisecond blinking to state transitions occurring over hours. We further demonstrate its capabilities in two biological systems. We performed cryo-SR-CLEM of YFP-labelled PML bodies in focused ion beam (FIB) milled lamellae from cultured mammalian cells. PML bodies are nuclear condensates that lack distinctive structural features for identification by cryo-ET alone, making cryo-SR-CLEM essential to localise them and resolve their internal organisation. We further show cryo-SR-CLEM in serial cryo-lift-out lamellae from *Nicotiana benthamiana* plant tissue expressing ATG9-eGFP, whose precise roles in autophagosome biogenesis remain incompletely resolved in plants. Together, these results establish VULCROM as a versatile platform for cryo-SR-CLEM and single-molecule cryo-photo-physics, addressing a critical gap in the instrumentation available for structural cell biology.

## Results

One major focus of our design concept for VULCROM (Fig. 1A,B) was to achieve high thermal and mechanical stability, accomplished through passive cooling and the high thermal capacity (~7,500 J/K) of the cryo-chamber. Passive cooling was implemented by thermally linking the cryo-chamber via a solid copper rod to a large LN_2_ reservoir (~12 L), eliminating the need for pumps and other sources of variations in the cooling rate. Cooldown requires ~50 min before samples can be loaded, with stable measurement temperature reached after ~90 min (details see suppl. Fig. S1). Measurements can be performed for ~12 h before the LN_2_ reservoir needs refilling. The high thermal capacity creates an inertia of the cryo-chamber regarding temperature changes, directly improving focus stability. To further improve mechanical stability, the objective lens (100× 0.9 NA, Nikon MUC11900) was mounted directly onto the copper structure inside the cryo-chamber, reducing the mechanical path between objective lens and sample stage for improved stability (for more details see suppl. Figs. S3, S4). Sample positioning and focusing are achieved by stacked piezo positioners inside the cryo-chamber (see Fig. 1B for design details). Together, these features enabled stability in the single-digit nanometre to angstrom-range over several hours (Fig. 1C-M).

**Figure 1.**
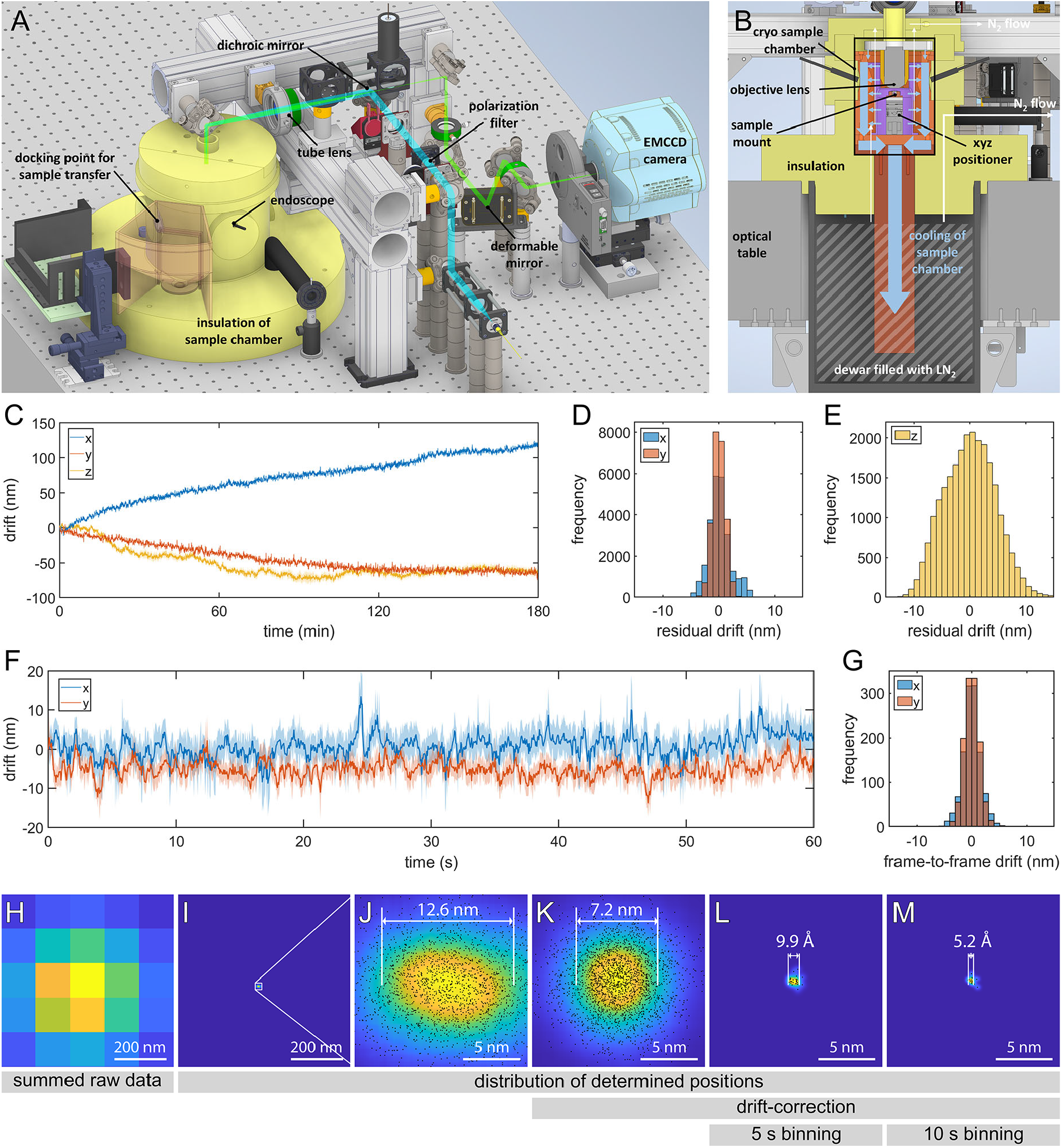
VULCORM setup and stability measurements. **A**: Schematic overview of VULCROM setup. The cryo-sample-chamber is housed within insulation material (yellow). A docking port allows sample transfer using the Leica Transfer Shuttle (insulation lid for transfer docking point not shown for visibility). Light paths for wide-field laser excitation and fluorescence detection are indicated by blue and green lines, respectively. **B**: Cut through cryo-sample-chamber of the VULCROM setup. Cooling (blue arrows) is achieved by a copper cup (orange) surrounding the interior of the cryo-sample-chamber and a copper rod that reaches into the LN_2_ reservoir underneath. White arrows indicate flow of N_2_ gas from LN_2_ boil-off through the sample-chamber to create a slight positive pressure to prevent build-up of ice contamination. Additional elements (purple) promote a laminar gas flow and guide the flow around the sample stage (grey). The objective lens is mounted inside the cryo-sample-chamber, heated to ambient temperature and insulated to protect it from the cold N_2_ gas atmosphere. An opening in the insulation of the cryo-sample-chamber on top of the heated back end of the objective lens allows guiding light in and out of the setup with the objective lens acting as a window. **C**: Drift measurement of 5 fluorescent microspheres over the course of 3 h (time intervals: 2 s). Solid lines represent the average drift, transparently coloured regions indicate the corresponding standard deviations. The drift is <1 nm/min. **D, E**: Residual drift was calculated by correcting each microsphere position by the drift determined with the remaining microspheres. FWHM of residual drift distributions in x-direction: 4.2 nm; in y-direction: 3.5 nm; in z-direction 11.3 nm. **F**: Drift measurement of 11 fluorescent microspheres over the course of 60 s with 50 ms time intervals. **G**: Histogram of frame-to-frame drift of measurement shown in **F** (FWHM: 3.2 nm in x-direction; 3.0 nm in y-direction). **H**: Summed image over 120 s of one of the microspheres used for drift measurements in **F** and **G.** Distributions of measured positions of the microsphere in **H** for every time interval are shown with individual localisation precisions in **I** (same scale) and **J** (magnified). Black points visualise individual positions (FWHM: 12.6 nm). **K**: Distribution of positions as in **J** after drift correction (FWHM: 7.2 nm). **L, M**: Distribution of positions after binning drift-corrected positions to 5 s intervals (**L**, FWHM: 9.9 Å) and 10 s intervals (**M**, FWHM: 5.2 Å).

The objective lens is fixed in a Macor plate that is kinematically mounted on top of the inner wall of the cryo-chamber to achieve high reproducibility when exchanging objective lenses. Set screws with ball tips in the plate allow angle adjustment. The Macor plate minimises thermal conductivity between the copper of the cryo-chamber and the objective lens. As the objective lens sits inside the cryo-chamber, it is heated to ambient temperature and thermally insulated to protect it from the cold N_2_ gas atmosphere (for more details see suppl. Fig. S2).

VULCROM utilises the N_2_ gas from LN_2_ boil-off to create a slight positive pressure inside the cryo-chamber (Fig. 1B and suppl. Fig. S3), minimising the build-up of ice contamination from atmospheric humidity. The N_2_ gas flow is regulated via a second gas outlet that releases boil-off gas directly from the LN_2_ dewar into the environment, and can be monitored via a temperature sensor at the N_2_ gas outlet of the cryo-chamber. Additional elements integrated into the cryo-chamber (Fig. 1B and Fig. S3, purple) promote laminar N_2_ gas flow around the sample stage, minimising vibrations.

To evaluate the stability of VULCROM, we performed measurements on fluorescent microspheres mounted on standard cryo-EM grids. Microsphere positions were determined by 2D Gaussian fitting, and axial coordinates were extracted by introducing astigmatism (1.0 µm RMS) via a deformable mirror in the detection path (see suppl. Fig. S5), enabling three-dimensional tracking. Figure 1C shows a representative stability measurement over 3 h, with a drift of <1 nm/min. Residual drift (Fig. 1D, E) was calculated according to Hoffman *et al*. ^11^ and represents the remaining localisation error after drift correction, setting the lower bound for achievable single-molecule localisation precision in the final cryo-SMLM image. The residual drift was 4.2 nm in x, 3.5 nm in y, and 11.3 nm in z (measured in FWHM of distributions of residual drift). These values correspond to the remaining error that would be present in a cryo-SMLM measurement after drift correction. As a result, every detected single-molecule position is not only scattered around the true molecular location due to its intrinsic localisation precision, but is also further dispersed by the position uncertainty of the remaining residual drift.

Where single-molecule events are not rare and sparsely distributed over time, the scattering of the residual drift averages out ^20^. This is illustrated in Fig. 1H-M, where the position of a single microsphere was tracked at 50 ms intervals over 120 s. The raw distribution of measured positions had a width of 12.6 nm (Fig. 1J), narrowing to an isotropic distribution with an FWHM of 7.2 nm after drift correction (Fig. 1K). Binning drift-corrected positions to 5 s and 10 s intervals further reduced this to 9.9 Å and 5.2 Å, respectively. We also evaluated stability on the millisecond timescale to assess whether fast vibrations or oscillations were present. Frame-to-frame drift was measured over 60 s at 50 ms time intervals (Fig. 1F), yielding a lateral movement FWHM of ~3 nm between subsequent frames (Fig. 1G).

We have integrated a deformable mirror (DM69-15, Alpao) in the detection path of VULCROM (see suppl. Fig. S5) to determine and correct optical aberrations ranging from subtle aberrations inherent to the optics of the setup to strong aberrations typical of imaging in thick specimens. We developed a sensorless adaptive optics (AO) correction routine with a tunable quality metric (see suppl. Fig. S6). This allows the routine to be optimised for high sensitivity to small aberrations such as PSF asymmetries at the nanometre level, or for broader features such as overall intensity and PSF width, making it adaptable to the range of scenarios encountered in cryo-FM imaging. AO correction of inherent optical aberrations of VULCORM is shown in suppl. Fig. S7.

To demonstrate the effect of aberrations caused by thick specimens on cryo-SR-FM, we imaged GFP-labelled endosomes in COS7 cells ~20 µm below the ice surface (Fig. 2). Internalised TRITC-labelled MSN-PEI nanoparticles, also present in the cells, were used to determine the optical aberrations via the AO correction routine (Fig. 2A-E), as their signal-to-noise ratio was substantially higher than that of the endosomes (SNR: ~18.5 vs ~1.5). AO correction improved wide-field image intensity by ~20% (Fig. 2F, G). The effect was more pronounced in cryogenic super-resolution optical fluctuation imaging (cryo-SOFI), where AO correction yielded a ~4-fold intensity increase (Fig. 2H-K), recovering weaker endosome signals that were absent without correction (compare Fig. 2J, K).

**Figure 2.**
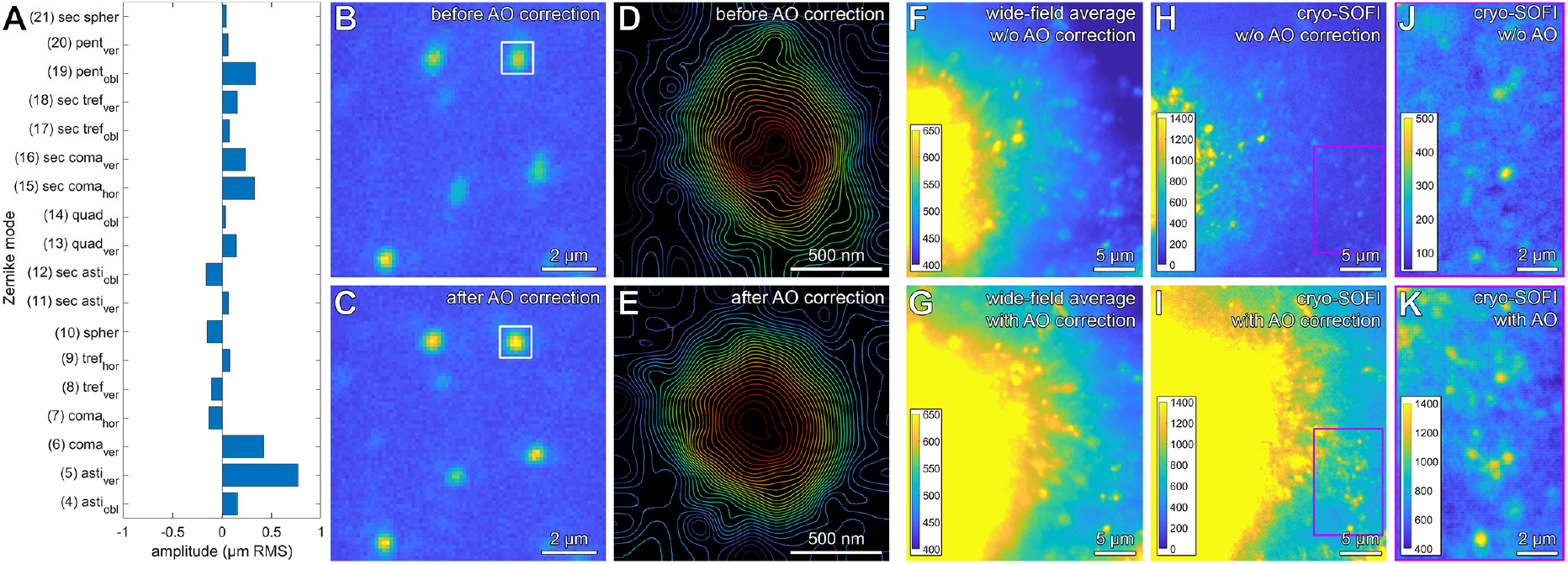
AO correction of aberrations for cryo-SOFI in thick specimens (GFP-labelled endosomes in COS7 cells ~20 µm below the ice surface). **A**: Amplitudes of Zernike modes determined to correct for optical aberrations caused by ~20 µm of ice, using TRITC-labelled MSN-PEI nanoparticles in COS7 cells. A qualitative comparison of cryo wide-field imaging before and after AO correction is given in **B** and **C**, respectively. **D** and **E** show contour plots of the PSF of one nanoparticle (indicated by white square in **B, C**) before (**D**) and after (**E**) AO correction. Intensity is displayed on a logarithmic scale. **F**-**K**: Cryo-FM imaging of GFP-labelled endosomes in the same COS7 cells, with and without AO correction. **F, G**: Comparison of wide-field averages with and without AO correction. Intensity gain with AO correction: ~20%. **H, I**: 2nd order cross-correlation cryo-SOFI of the same area as shown in **F** and **G** with and without AO correction. Intensity gain with AO correction: ~4-fold. **J, K**: Magnified images of areas marked with the purple rectangle in **H** and **I** with different intensity scaling to achieve a similar dynamic range.

To demonstrate the capabilities of VULCROM for studying single-molecule cryo-photo-physics, we performed measurements of single Atto647N organic dye molecules in vitreous ice over ~2.5 h. Such measurements require high mechanical stability both over the course of hours, to avoid out-of-focus drift, and at the millisecond timescale, to avoid interference from fast vibrations with the recorded single-molecule intensity profiles. Data were acquired at 20 ms camera integration time and a laser intensity of 50 W/cm^2^. Before acquisition, two images at lower laser intensity (5 W/cm^2^) were recorded with perpendicularly oriented linearly polarised excitation light (Fig. 3B), exploiting the fixed dipole moments of fluorophores under cryo-conditions ^3, 4, 5, 21^ to distinguish isolated single molecules from clusters. Figure 3C shows time profiles of the full ~2.5 h measurement binned to 2 s, alongside selected intervals binned to 20 ms, capturing photo-blinking across timescales from milliseconds to hours. Molecule 1 showed frequent on-peaks during the first 30 min, remained off for over an hour, then resumed switching until the end of the measurement. Molecule 2 remained in an on-state for the entire ~2.5 h. Position 3, identified as multiple fluorophores from the polarisation data, showed distinct on-peaks throughout, with on-times of a few seconds and off-times of 10-100 s, and intensity values several times larger than those of the single isolated molecules. At 20 ms temporal resolution, fast millisecond blinking was observed within all on-states, including the sharp intensity peaks of positions 1 and 3 (Fig. 3C), suggesting a potential relationship between fast millisecond cryo-photo-blinking and the longer-lasting on- and off-state transitions occurring over seconds to hours.

**Figure 3.**
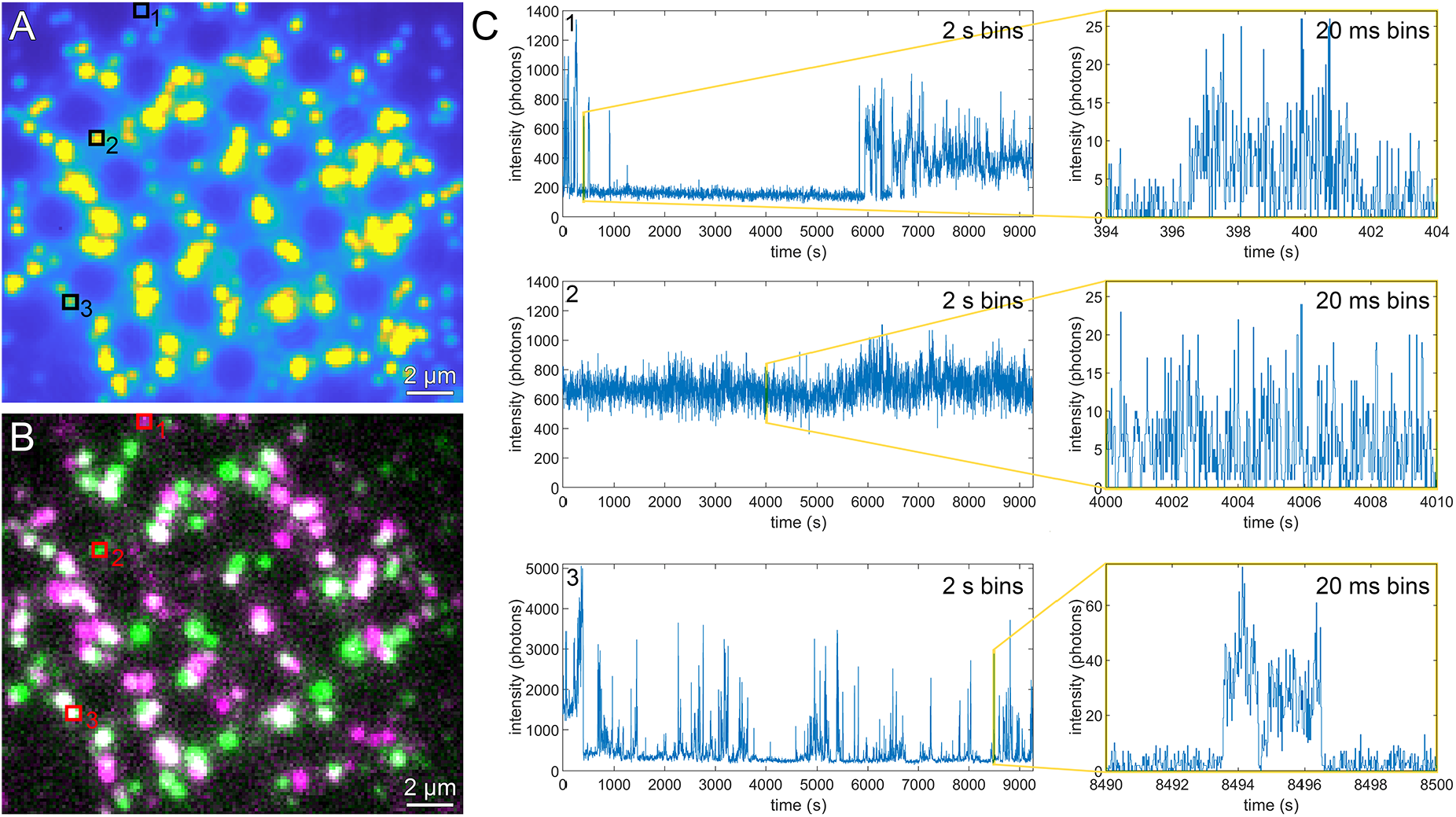
Single-molecule blinking of Atto647N dyes under cryo-conditions on different time scales. **A**: Summed image of time series acquired with circularly polarised excitation light at a laser intensity of 50 W/cm^2^. **B**: Overlay of two fluorescence images recorded before the acquisition of the time series with linearly polarised excitation light at two perpendicular orientations (laser intensity: 5 W/cm^2^). Magenta or green spots indicate high probability of isolated single molecules; white spots indicate high probability of multiple molecules within a diffraction limited area. **C**: Example time series of single molecules (1 and 2) and multiple molecules in close proximity (3) at the positions indicated by black squares in **A.** The left time series show the entire recording of 2.5 h and are binned to 2 s. The right-hand time series show 20 ms bins of the short intervals highlighted by yellow lines in the full recordings. All on-states in the 2 s binned time series show fast blinking on the millisecond time scale.

We designed a docking port compatible with the Leica Transfer Shuttle (see suppl. Fig. S4 for sample temperature during transfer), enabling direct integration of VULCROM into standard cryo-CLEM workflows. Compatibility with Leica Autogrid cartridges allows samples to be transferred between VULCROM and a cryo-FIB milling instrument while maintaining grid orientation, supporting stepwise iterative milling workflows with cryo-SR-FM imaging between milling steps.

To demonstrate cryo-SR-CLEM with VULCROM, we have imaged two distinct biological systems (Figs. 4 and 5). Cryo-SMLM of YFP-labelled promyelocytic leukemia protein (PML) bodies in FIB-milled lamellae from human foreskin fibroblast (HFF) cells show the arrangement of PML-YFP within PML bodies in the structural context provided by cryo-ET (Fig. 4). Cryo-SMLM data was acquired at 488 nm and 200 W/cm^2^ and resolved the PML-YFP with an average single-molecule localisation precision of 7.7 nm (suppl. Fig. S8), revealing a distinct PML-YFP shell surrounding a PML-free core. Relaxing the filter criteria to include weaker single-molecule signals increased the number of detected molecules by more than 3-fold, at an average localisation precision of 12.6 nm (suppl. Fig. S8), illustrating how cryo-SMLM can be tuned to the needs of a given application. Precise fluorescence-EM correlation was essential for identifying PML bodies in the cryo-ET data, as these nuclear condensates appear as featureless smooth-textured spherical regions (400-800 nm diameter) that cannot be identified by cryo-ET alone (Fig. 4D, E). Full molecular characterisation of PML body architecture will be presented in a forthcoming publication.

**Figure 4.**
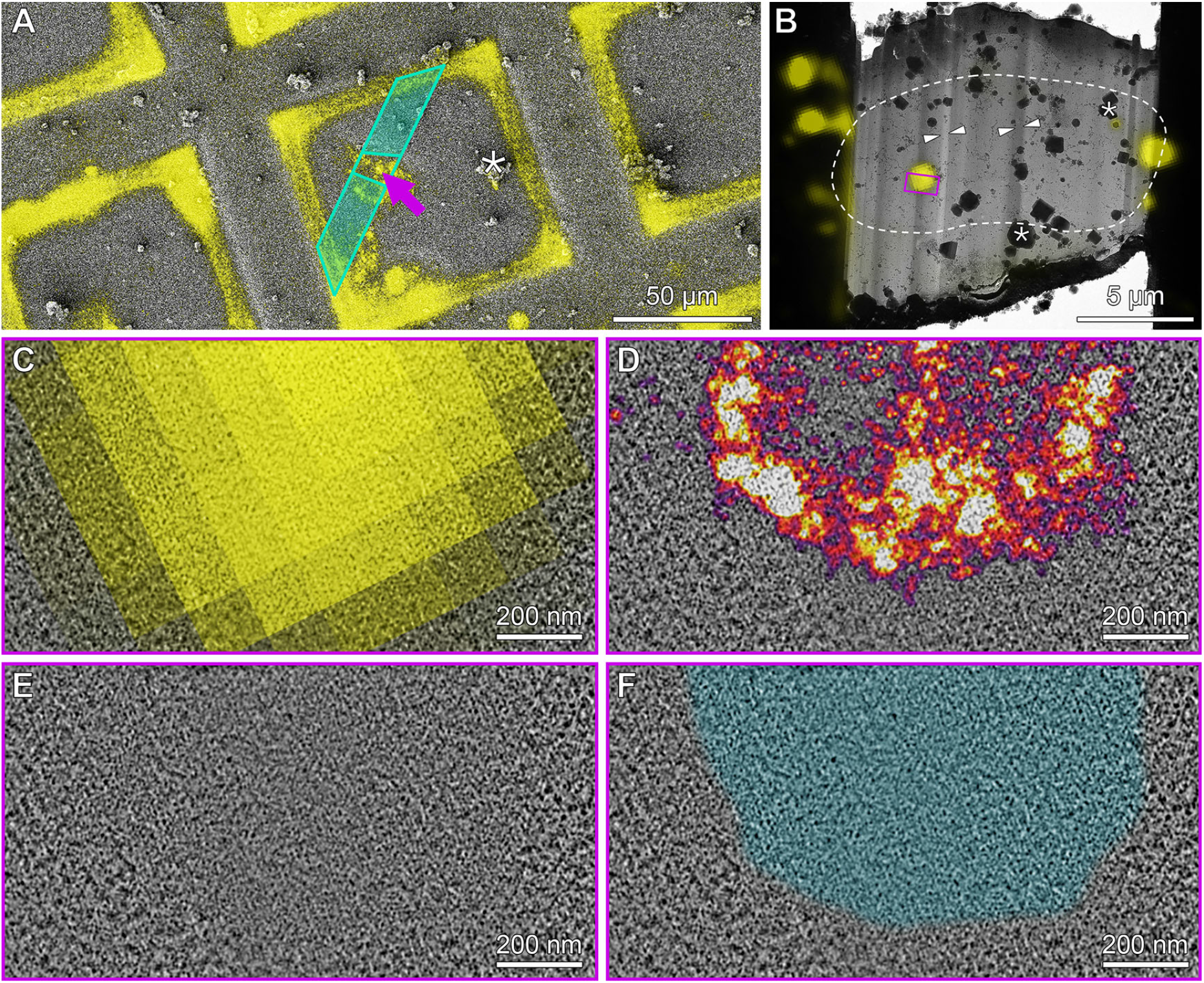
Cryo-SR-CLEM of YFP-labelled PML bodies in the nucleus of HFF cells. **A**: Overlay of cryo-SEM image and in-column cryo-fluorescence used to identify candidate cells for FIB-milled cryo-lamella preparation **B**: TEM micrograph of a cryo-lamella overlaid with conventional cryo-FM signal. Magenta box indicates the area shown in **C.** Dashed white line indicates the approximate outline of the nucleus. White arrowheads indicate stripes of uneven lamella thickness. White asterisks indicate crystalline ice deposition on the lamella surface introduced during the three specimen transfers between microscopes. **C**: Slice of a tomogram (53,000× nominal magnification) and overlay with conventional cryo-FM image of area highlighted with purple rectangle in **B** and arrow in **A. D**: Overlay of corresponding cryo-SMLM image (acquired with 488 nm laser at 200 W/cm^2^) of the same area as in **C. E**,**F**: Corresponding tomogram slice without fluorescence overlay (**E**) and with manually segmented (blue) PML body (**F**).

**Figure 5.**
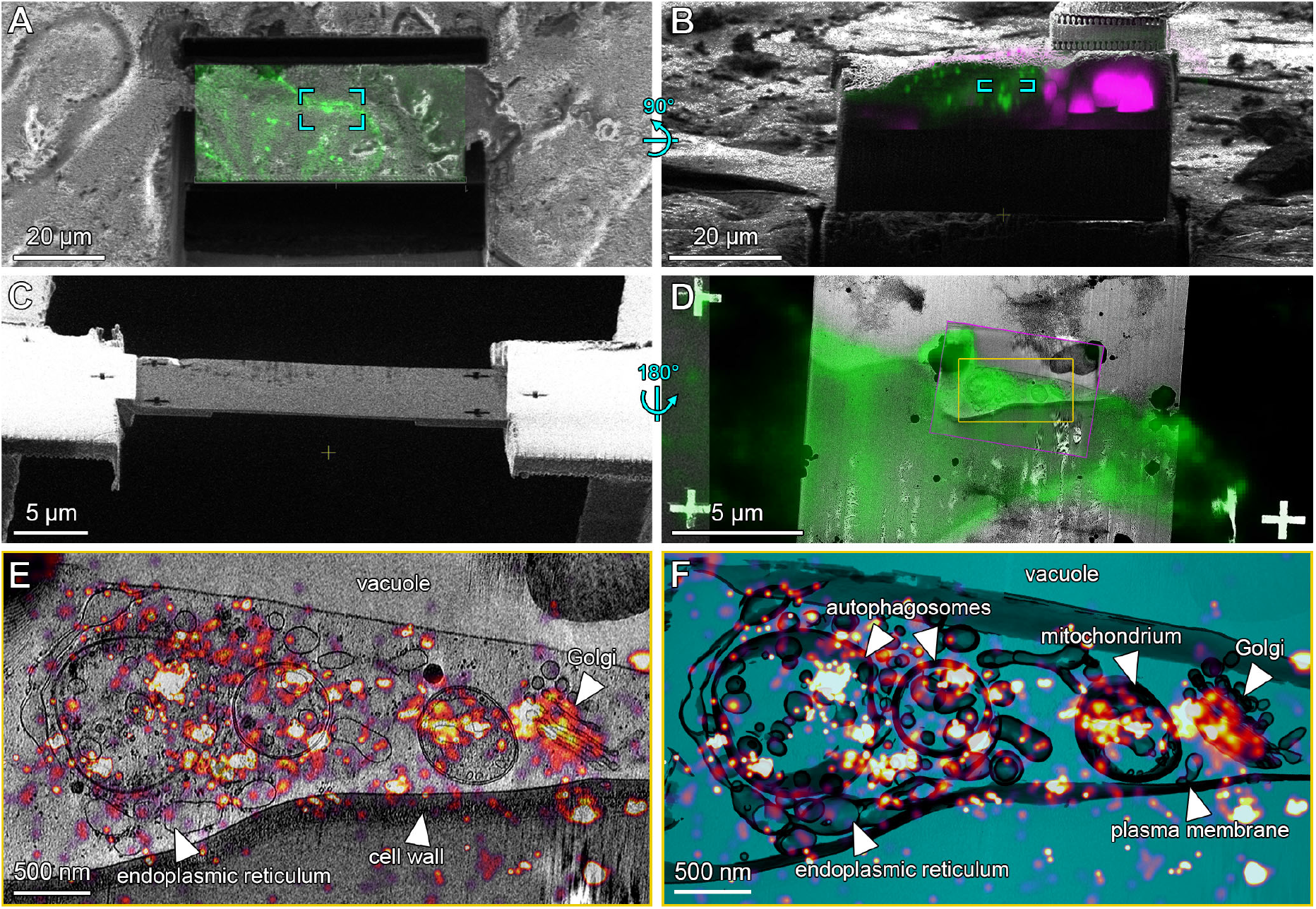
Cryo-SR-CLEM of *N. benthamiana* leaf expressing ATG9-eGFP. **A**: Cryo-SEM overview of a high-pressure frozen *N. benthamiana* leaf where a ~50 µm × 20 µm × 20 µm block was milled using FIB in preparation for cryo-lift out. Maximum intensity projection through 15 µm deep confocal cryo-FM volume was overlaid. Cyan box indicates the location of a polished lamella shown in **D. B**: The milled block was attached to an EasyLift (Thermo Fisher Scientific) needle and lifted out. Shown is an image of the block at a shallow angle (10°) overlaid with a maximum intensity projection of the confocal volume. Chloroplast fluorescence is shown in magenta. **C**: The block was moved into an empty electron microscopy grid and sectioned into ~ 2 µm sections that were subsequently thinned to ~ 150 nm. Shown is the final polished lamella at a shallow angle. **D**: TEM overview (2250× nominal magnification) of the cryo-lamella overlaid with conventional cryo-FM. **E**: TEM micrograph (8500× nominal magnification, purple rectangle in **D**) and overlay with cryo-SMLM image (acquired with 488 nm laser at 200 W/cm^2^) of area marked by yellow rectangle in **D** showing two autophagosomes, a mitochondrion and a Golgi (left to right). **F**: Overlay of segmented membranes with cryo-SMLM image of the same area as in **E**.

To further demonstrate the versatility of VULCROM in the context of structural cell biology, we performed a proof-of-concept demonstration of cryo-SR-CLEM on a serial cryo-lift-out lamella of *Nicotiana benthamiana* leaf expressing ATG9-eGFP, to investigate the nanoscale organisation of ATG9 (Fig. 5). ATG9 is a key factor in plant host immunity, yet its subcellular distribution remains poorly understood. The serial lift-out approach ^22^ was used to obtain a thin section of high-pressure frozen *N. benthamiana* leaf tissue, which were then thinned to a ~150 nm thick lamella using FIB. Cryo-SMLM data were acquired using a 488 nm laser at 200 W/cm^2^, enabling localisation of single eGFP-labelled ATG9 molecules with an average precision of 21.1 nm. Cryo-SR-CLEM enhanced the spatial resolution of ATG9-eGFP, making it possible to assign its signal to specific compartments of the segmented cryo-ET volume (Fig. 5F). Consistent with current understanding, strong labelling was observed at the Golgi apparatus and at two putative autophagosomes. More specifically, most localisations were associated with ~100 nm vesicles located either within, or in close proximity to the autophagosomes, and the cis face of the Golgi. In contrast, very few molecules were detected near the putative endoplasmic reticulum, despite its close association with this organelle complex. Additionally, dense fluorescent labelling was observed around inner mitochondrial membranes in the same region, although the significance of this remains unclear. These observations demonstrate that cryo-SR-CLEM can resolve heterogeneous ATG9 localisation both within and between organelles, generating structural hypotheses for further investigation.

## Discussion

VULCROM is a setup that can be used for various applications ranging from detailed single-molecule cryo-photo-physics analysis to cryo-SR-CLEM in vitreous biological specimens. We have shown that this design minimises drift and vibrations to well below 10 nm for cryo-SMLM measurements over several hours (Fig. 1C-E). For applications not based on rare single-molecule events scattered over long time periods, VULCROM offers stability and localisation precision down to the angstrom-range (Fig. 1H-M). This is the same level as what has so far only been achieved with vacuum-insulated cryostats ^13, 19^. This high stability is primarily achieved through passive cooling, the large thermal capacity of the cryo-chamber, and precise heating of the objective lens, minimising temperature variations and ensuring high stability from milliseconds to hours (Fig. 1F, G). Investigating single-molecule cryo-photo-physics requires stability over long timescales. Without this, vibrations or other movements in the setup, especially those occurring on comparable timescales to the molecular dynamics, can introduce artefacts and lead to misinterpretation. The example measurements shown in Fig. 3 illustrate the importance of setup stability from the millisecond to the hours range. The single-molecule data indicates that there might be a relationship between fast cryo-photo-blinking on the milliseconds time scale and long-lasting on- and off-states in the range of seconds to hours.

Our setup achieves sub-10-nm resolution with cryo-SR-CLEM in vitreous biological specimens, including full compatibility with cryo-ET workflows. To validate the overall performance of the VULCROM setup for correlative imaging, we have performed cryo-SR-CLEM experiments of different biological application examples (Fig. 4-5). These examples show that with utilizing cryo-SMLM we can achieve a correlation of fluorescence information with structural information of cryo-ET that is far beyond the capabilities of conventional cryo-CLEM. We showed that cryo-SR-CLEM can be successfully performed even for challenging cryo-ET workflows such as serial cryo-lift-out procedures (Fig. 5; see suppl. Fig. S10 for more details regarding our cryo-SR-FM workflow). The strength of cryo-SR-CLEM lies in visualising fluorescently labelled features in cryo-tomograms that would otherwise be difficult or impossible to identify. This is well illustrated by our cryo-SR-FM and cryo-ET of PML bodies. While cryo-ET reveals a difference in texture between PML bodies and the surrounding chromatin, confident identification of the smooth regions containing both shell and core without cryo-SR-FM would not be possible. Furthermore, the distribution of PML-YFP within the body could not be resolved by cryo-ET alone given the small size of the protein in the crowded cellular environment.

For cryo-SMLM applications, the stability of the setup defines the lower boundary of achievable single-molecule localisation precision and resolution. The average localisation precision across the biological applications shown here ranged from 7 to 21 nm, reflecting contributions from both sample and data processing factors. Sample-related factors include fluorophore photo-physics, background fluorescence, and specimen thickness. Camera integration time and temporal binning directly influence localisation precision; ideally, integration time should match the average fluorophore on-time to maximise SNR, though longer integration times also increase background. The SMLM filter criteria allow tuning between maximum localisation precision and higher molecular detection density (for example see suppl. Fig. S8), and can be adjusted to suit the needs of a given application.

The deformable mirror integrated into the VULCROM detection path serves multiple purposes. It enables measurement and correction of residual aberrations inherent to the setup itself (suppl. Fig. S7), arising for example from the temperature gradient across the objective lens or refractive index mismatch caused by the cold N_2_ gas between the objective front lens and the specimen. It can also introduce astigmatism or other PSF deformations to enable axial localisation of single molecules or small fluorescent objects (Fig. 1C). The most powerful application of adaptive optics in cryo-SR-FM, however, is the correction of aberrations caused by thick specimens, where image quality deteriorates rapidly and severely impacts super-resolution imaging even with robust methods such as cryo-SOFI (Fig. 2H-K). Aberration correction will become increasingly important in cryo-CLEM in general as cryo-ET in tissues and organoids gains popularity through cryo-lift-out techniques ^22, 23, 24^. Cryo lift-out can be used to obtain specimens thin enough for cryo-ET even in cases where direct lamella milling is not possible due to the sample thickness. Here, high-precision 3D cryo-CLEM is required for targeting rare objects and events. This is an approach that is both technically challenging and instrument time-consuming, meaning that high quality cryo-FM data is crucial for choosing optimal specimen areas. AO will be important to correct the aberrated PSF in thick specimens both to improve the image quality, but also to minimise the error in position determination.

Our design concept reduces complexity and costs compared to vacuum-insulated systems. Vacuum insulated systems offer, in principle, lower levels of crystalline ice deposition during imaging, at the cost of flexibility, modularity, and integration with existing cryo-ET workflows. For example, the sample transfer of VULCROM can be adjusted to fit individual requirements (e.g. compatibility with sample cartridges and transfer systems with FIB or TEM). The sample transfer mechanism described here is compatible with commercial cryo-CLEM equipment, which allows to integrate cryo-SR-FM into various different cryo-CLEM and cryo-ET workflows. Here we have demonstrated the feasibility of cryo-ET imaging directly after cryo-FM, which is the optimal approach where instrument time is limiting. The possibility of going back and forth between FIB-milling and cryo-SR-FM imaging enables iterative milling and imaging workflows, similar to as shown by Wu *et al*. ^25^ for conventional cryo-CLEM. Additionally, it also allows utilizing the FIB for removing ice contamination gathered during the sample transfer steps before cryo-ET data acquisition. The vacuum-free design of the VULCROM system furthermore enables the implementation of cryo-immersion imaging ^26, 27, 28^ or alternative objective lens geometries (e.g. dual-objective lenses ^29^ or 4Pi configuration ^30^), which would allow to further increase resolution and sensitivity.

We have demonstrated the performance of VULROM for applications ranging from single-molecule cryo-photo-physis experiments to cryo-SR-CLEM in vitreous biological specimens. Combined with its flexible and modular design concept, VULCROM has the potential to establish cryo-SR-CLEM as a powerful and broadly accessible tool for structural cell biology.

## Materials and Methods

### VUCROM setup

An MLE LightHUB+ system (Omicron) with lasers of the wavelengths 405 nm (LuxX 405-120, Omicron), 488 nm (LuxX 488-200, Omicron), 561 nm (OBIS 561-100, Coherent) and 647 nm (LuxX547-140, Omicron) coupled into a single-mode fibre was used as an illumination source. The laser light was collimated using a lens with 100 mm focal length (AC254-100-A-ML, Thorlabs). The illuminated area can be changed using an iris diaphragm (SM1D12D, Thorlabs). Further, to switch between the maximum size and cropped size of the illuminated spot, a two-position slider (ELL6B, Thorlabs; one position with iris diaphragm and other position empty) was installed in the excitation pathway. Another two-position slider mounted with a custom-made 45°-mirror mount was utilised to guide the light of a white LED (LEDW7E, Thorlabs) into the pathway, which was used for reflection imaging. A quarter wave plate (10RP54-1B, Newport) mounted on a kinematic flip mount (Kin-a-Flip, Newport) was used to convert the elliptical polarisation of the laser to two orthogonal liner polarisations if needed. The polarisation was measured after the objective lens using a film polariser (LPNIRE100-B, Thorlabs) and a power meter (S121C and PM100D2, Thorlabs). The collimated light is reflected from a quad-line dichroic mirror (zt405/488/561/647rpc, Chroma) into the tube lens (focal length 200 mm, MXA20696, Nikon), which focuses the light into the back focal plane of the objective lens (CFI TU Plan Apo EPI 100×, N.A. 0.90, working distance 2.00 mm, MUC11900, Nikon) for a wide-field illumination. The fluorescence light of the sample is collected by the objective lens and focused by the tube lens and two relay lenses (67331, Edmund Optics) with a focal length of 160 mm each onto an electron multiplying CCD (EMCCD) camera (iXon Ultra 888, Andor). The fluorescence light is passed through the dichroic mirror and a specific bandpass filter (details see Material and Methods for individual experiments) mounted in a motorised filter wheel (FW102C, Thorlabs). A deformable mirror (DM69-15), was placed at the conjugate plane of the back focal plane of the objective lens between the two relay lenses in the detection path (for more details see Fig. S5).

### Preparation of fluorescent spheres samples and parameters for drift measurements

The stock solution of yellow-green 0.17 µm FluoSpheres (Thermo Fisher Scientific) was diluted 1:10 in ethanol and sonicated for 1 minute. EM-grids (C-Flat 200 Mesh, CF-2/1-Cu-50, 2/1, Protochips) were used as sample carrier and the grids were glow-discharged with high RF level setting in a plasma cleaner (Harrick Plasma) for one minute. A drop of ~3 µl of the diluted FluosSpheres was pipetted on a grid and after the solvent has evaporated, the gids were clipped into an AutoGrid ring (Thermo Fisher Scientific). For the drift measurements the signal of multiple fluorescent microspheres was recorded for up to 180 min with an integration time of the EMCCD camera of 10-20 ms and an EM gain of 50-300. The 488 nm laser was set to an intensity of 15-20 W/cm^2^ for fluorescence excitation. For 3D-drift measurements, vertical astigmatism of 1.0 µm RMS was introduced in the detection by using the deformable mirror.

### Single-molecule cryo-photo-physics sample preparation and measurements

ATTO647N (ATTO-TEC) molecules were diluted to a concentration of 0.01 ng/µl in DPBS and sonicated for a few minutes. The same EM-grids and preparation procedure as in the preparation of the fluorescent microspheres was used. A 3,5 µl droplet of this dilution was pipette onto an EM-grid and blotted with a filter paper for approx. one second. Directly afterwards the sample was plunged into liquid ethane-propane using a manual plunge-freezer, clipped into an AutoGrid ring (Thermo Fisher Scientific) and safely stored in a grid box in a liquid nitrogen storage dewar. Grids with vitrified dye solution were transferred into the VULCROM setup with a Cryo Transfer Shuttle (Leica Microsystems). To acquire polarisation information of the molecules, an image of each excitation polarisation was acquired for four minutes (accumulation of 1,200 frames with 200 ms integration time). The laser spot of the 647 nm laser was adjusted to diameter of ~20 µm with an intensity of 20 W/cm^2^. For imaging a band pass filter (700 nm/75 nm, Chroma) was used. The electron multiplying gain of the camera was set to 300. For the cryo-photo-physics measurement the laser intensity was increased to 50 W/cm^2^ and the electron multiplying gain of the camera to 700. Data was recorded for 154 min in total (462,000 frames with an integration time of 20 ms).

### Sample preparation of COS7 cells with fluorescently labelled endosomes and nanoparticles

COS-7 cells were cultured in Dulbecco’s Modified Eagle Medium (DMEM) (Sigma-Aldrich) supplemented with 10% (v/v) fetal bovine serum (FBS) (Sigma-Aldrich, USA), 2% (v/v) GlutaMAX supplement (Gibco, Thermo Fisher Scientific), and 1% (v/v) sodium pyruvate (Sigma-Aldrich). Cells were seeded in 24-well plates and labelled with CellLight Early Endosomes-GFP (BacMam 2.0, Thermo Fisher Scientific). After labelling, cells were incubated overnight to allow expression of the GFP marker. On the following day, labelled cells were transferred onto EM-grids (Ultrafoil, R2.2. mesh 200, Quantifoil) for nanoparticle incubation. Surface hydrophilisation of grids was performed by glow-discharge treatment using a plasma cleaner (GloQube, Quorum). Glow-discharged grids were transferred into an Ibidi 2-well co-culture dish, with 1–2 grids placed per well. Each well was incubated with 10 µL fibronectin (50 µg/mL) solution at 37 °C for 2 h. COS-7 cells with GFP-labelled endosomes were detached using trypsin-EDTA and incubated at 37 °C for 3 min. Trypsin activity was neutralised by the addition of complete culture medium. Cells were collected by centrifugation at 300 g for 3 min and resuspended in 1 mL of complete medium. For seeding, 5 µL of the cell suspension (approximately 8 × 10^3^ cells) was added to each well containing fibronectin-coated EM-grids. Cells were allowed to attach and spread on the EM-grids overnight. TRITC (tetramethylrhodamine isothiocyanate)-labelled mesoporous silica nanoparticles at concentration of 10 µg/mL were added after sonification to the endosome-labelled COS-7 cells. Cells were incubated with nanoparticles for 2 h at 37 °C in a humidified atmosphere containing 5% CO_2_ prior to plunge freezing. EM-grids containing adherent COS-7 cells were vitrified using a manual plunge-freezer and mounted into AutoGrid cartridges.

### Sample preparation of PML bodies

Human foreskin fibroblasts (HFF) stably expressing PML1-YFP under doxycycline-inducible control were cultured as previously described ^31^. Cells were initially maintained with 5 μg/ml hygromycin (Invitrogen, 10687-010) and 1 μg/ml puromycin (Sigma-Aldrich, P8833) for hTERT expression and selection, respectively. During experiments, HFF cells were cultured in Minimum Essential Medium Eagle (Sigma-Aldrich) supplemented with 10% fetal bovine serum, 2 mM L-glutamine (Life Technologies, 25030-024), 1 mM sodium pyruvate (Life Technologies, 11360-039), and 100 U/ml penicillin/streptomycin. For cryo-EM sample preparation, Quantifoil SiO2 R2/1 200-mesh gold grids were treated with laminin for 30-40 minutes and air-dried on Whatman 40 filter paper. Grids were placed in MatTek glass-bottom dishes containing 2 ml culture media. HFs were seeded onto grids and cultured for 12-14 hours to achieve optimal confluence for FIB milling. PML1-YFP expression was induced by doxycycline treatment for 7 hours prior to vitrification. Vitrification was performed using a Vitrobot Mark IV (Thermo Fisher Scientific). To ensure optimal structural preservation, excess medium was manually blotted from the grid backside using Whatman 40 filter paper 2 minutes before freezing. Subsequently, 3 μl of 8-9% glycerol in medium was applied to the cell-bearing side as a cryoprotectant. Grids were immediately plunge-frozen in liquid ethane and stored in liquid nitrogen until FIB-milling.

### Sample preparation of *N. Benthamiana*

The ATG9:eGFP construct was generated using Gibson Assembly, following the protocol described previously ^32, 33^. Specifically, the vector backbone was a pK7WGF2 derivative domesticated for Gibson Assembly, containing a C-terminal fluorescent eGFP tag. Wild-type *Nicotiana benthamiana* plants were grown in a controlled-environment chamber at 24 °C. Plants were cultivated in a substrate composed of organic soil mixed with sand and vermiculite at a 3:1 ratio (Levington’s F2 with sand and Sinclair’s 2–5 mm vermiculite). Growth conditions consisted of high-intensity light under a long-day photoperiod of 16 h light and 8 h dark. All experiments were performed using 4-week-old plants. Agrobacterium-mediated transient expression of ATG9:eGFP in *N. benthamiana* was performed via leaf agroinfiltration as previously described ^34^. *Agrobacterium tumefaciens strains* carrying the ATG9:eGFP construct were rinsed with water and resuspended in infiltration buffer (10 mM MES, 10 mM MgCl_2_, pH 5.7). The optical density at 600 nm was measured using a BioPhotometer spectrophotometer (Eppendorf). The final suspension was infiltrated into the leaves of 4-week-old *N. benthamiana* plants using a needleless 1 mL Plastipak syringe.

### High pressure freezing

*N. benthamiana* leaves were frozen 3 days post infiltration with *A. tumafeciens*. They were cut into ~1 cm × 2 cm sections and vacuum infiltrated using a (washed and reused) plastic syringe with cryoprotectant buffer (40% Ficoll 70 (Sigma), 180 mM glycerol and 20 mM MES pH 5.5). A 2 mm diameter biopsy punch was used to excise sections. These were placed in the 300 µm recess of 3 mm diameter type-B carrier submerged in cryoprotectant buffer and closed with the flat side of another type-B carrier (Wohlwend). Freezing was done using HPM010 (Baltec).

### Correlative serial lift-out

Frozen carriers were first screened using confocal cryo-FM to locate potential regions of interest. Planchettes were mounted on Autogrids (Thermo Fisher Scientific) were imaged using Stellaris 5 (Leica) confocal microscope equipped with a cryogenically cooled stage. Confocal Z-stacks were acquired at 108 nm pixel size and 700 nm Z step. These planchettes were then brushed thoroughly using a painter’s brush to remove accumulated ice contamination and loaded into Aquilos 2 (Thermo Fisher Scientific). Image correlation between cryo-FM and SEM signal was done by making use of the unique pattern of cells on the leaf surface. Serial lift-out was then performed as described previously ^34^. An approximately 50 µm × 25 µm × 20 µm block was cut using FIB-milling, attached to a lift-out needle and transferred to a 400 × 100 mesh lift-in grid (Agar Scientific). The block sectioned into ~2 µm thick blocks; these were checked using an integrated fluorescence microscope (iFLM, Thermo Fisher Scientific) to identify sections containing fluorescent signal. The FIB-facing face of these blocks was polished before coating with ~200 nm thick protective organoplatinum layer. Lamella were then cut to ~200 nm final thickness.

### Focused ion beam milling of PML bodies

Grids were clipped in Autogrids and loaded into Aquilos 2. The iFLM integrated microscope was used to identify cells with promising fluorescence and correlated with SEM signal using Maps (Thermo Fisher Scientific). Lamella were cut using AutoTEM (Thermo Fisher Scientific) and polished manually to target thickness of ~ 200 nm.

### Cryo-EM and cryo-ET data acquisition

Grids were loaded into a Titan Krios (Thermo Fisher Scientific) equipped with K3 and BioQuantum energy filter (Gatan). Tilt series were acquired using SerialEM and PACE tomo ^35, 36^ at 3°, aiming for −66° to 66° tilt range. A 70 µm aperture and 20 eV slit was used during data acquisition. Tilt-series for tomogram shown in Fig. 5 (*N. benthamiana* leaf) was acquired at 0.45 e^−^ per frame for a total fluence of 20.4 e^−^Å^−1^. Tilt-series for tomogram shown in Fig. 4 (PML) was acquired at 4.5 e^−^Å^−1^ per tilt fractionated into 7 frames, resulting in a total fluence of 187.4 e^−^Å^−1^.

### Cryo-ET data processing

Frames were aligned using Alignframes (IMOD); tilt series were aligned and tomograms reconstructed using Etomo ^37^. 3dmod (IMOD) and Fiji were used for tomogram visualisation. The tomogram of *N. benthamiana* leaf tissue was segmented using MemBrain. False positive membrane assignment was removed using the volume eraser tool in Chimera 1.9 ^38^ and rotated to match the orientation of the untilted specimen before projection.

### AO and cryo-SOFI measurements

AO correction routine was performed using TRITC-labelled MSN-PEI nanoparticles in COS7 cells under an ice layer of ~20 µm. A laser intensity of 20 W/cm^2^ of the 561 nm laser. Individual images were captured with an integration time of the camera of 400 ms and an electron multiplication gain of 180. For ever image a different amplitudes of a certain Zernike mode (ranging from primary astigmatism to secondary spherical aberration) was applied in an iterative fashion following the sensorless correction routine illustrated in suppl. Fig. S6. After 20 iterations over all Zernike modes those amplitudes were selected that resulted in the best value of the quality metric. This was used for subsequent AO correction of the GFP-labelled endosomes in the same cell. For cryo-SOFI measurements of the GFP-labelled endosomes a laser intensity of 75 W/cm^2^ of the 488 nm laser was used for fluorescence excitation and stimulation of fluorescence intensity fluctuations. Raw data was recorded with an exposure time of 50 ms and an electron multiplying gain of 200 over the course of 600 s. The data set without AO correction was recorded prior to the data set with AO correction (both with same parameters). Data was binned to 200 ms to increase the signal-to-noise ratio for the cryo-SOFI reconstruction. Cryo-SOFI reconstruction was done as described previously^15^.

### Cryo-SMLM data acquisition

A laser intensity of 200 W/cm^2^ of the 488 nm laser was used for fluorescence excitation and stimulation of photo-blinking. The diameter of the illuminated area in the sample plane was adjusted to a value smaller than the width of the lamella. This prevents heating of the surrounding area, which absorbs substantially more light due to the thickness of the un-milled specimen and presence of the support film. Raw data was recorded with an exposure time of 50 ms and an electron multiplying gain of 140 (YFP-PML) or 700 (eGFP-ATG9) over the course 2.6 h (YFP-PML) or 1.4 h (eGFP-ATG9). Data was binned to 500 ms to increase the signal-to-noise ratio for the cryo-SMLM reconstruction. Cryo-SMLM reconstructions were done as described previously ^10^.

### Correlation of cryo-FM and cryo-EM/ET images

Transformation of coordinate systems for correlation of cryo-FM with cryo-EM/ET was done as described previously ^15^. Specifically, features identifiable in reflected light or fluorescence imaging and in cryo-EM/ET were used as reference positions. These were either distinct features of the outline of the lamella and ice contamination on the lamella (PML bodies) or crosses that have been milled into the lamella (ATG9). We used the Control Point Selection Tool of MATLAB (Mathworks) to determine the coordinate transformation between the fluorescence and the cryo-EM/ET based on the identified reference positions. The same approach was used to correlate cryo-EM/ET images of different magnifications.

## Acknowledgements

The authors gratefully acknowledge Professor Jessica Rosenholm, Åbo Akademi University, Finland, for providing mesoporous silica nanoparticles. We thank Chris Boutell, Centre for Virus Research, Glasgow, UK, for generously supplying the human foreskin fibroblasts stably expressing PML1-YFP. This work was supported by a Volkswagen Foundation Freigeist Fellowship to R.K. (91671; 91671-2); by the Deutsche Forschungsgemeinschaft (DFG) via grant No. 496128632 to R.K.; by a UK Research and Innovation Future Leaders Fellowship (MR/W010690/1) to S.D.C.; a Medical Research Council programme grant (MR/Z504464/1) to S.D.C. and a Biotechnology and Biological Sciences Research Council (BBSRC) grant (BB/X016382/1) to T.O.B. We thank the Advanced Light and Fluorescence Microscopy Facility (ALFM) at CSSB for support with light microscopy image acquisition. All electron microscopy was performed at the Multi-User CryoEM Facility at CSSB, supported by the University of Hamburg and DFG grant numbers INST 152/772-1|152/774-1|152/776-1 FUGGand.

## Author contributions

R.K. conceived and supervised the project. J.F., V.Q.D., N.P. and R.K. designed and built the VULCROM setup and implemented algorithms and software for associated hardware control and data analysis. J.F., N.P., I.H., E.L.H.Y., T.O.B., S.D.C. and V.P. prepared and provided samples. J.F., N.P., V.P. and R.K. performed measurements and data analysis. All authors contributed to the manuscript writing and editing.

## Data and code availability

CAD drawings (Autodesk Inventor files) for the VULCROM setup can be downloaded via the following link: https://github.com/rainerkaufmann/VULCROM. MATLAB (Mathworks) code for the sensorless adaptive optics correction routine for cryo-FM can be downloaded via the following link:https://github.com/rainerkaufmann/cryoAO. Source data will be made available upon acceptance.

