## Supplementary Figures for "On-lamella super-resolution cryo-CLEM for cryo-ET enabled by vacuum-free ultra-stable cryogenic fluorescence microscopy"

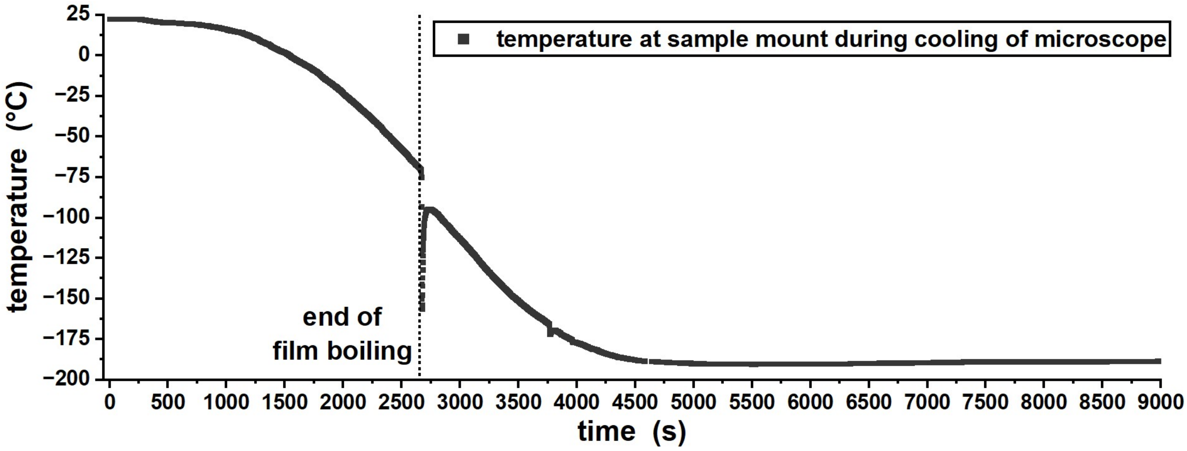


**Figure S1 | Temperature at sample mount during cooldown of the cryo-sample-chamber.** The course of the temperature was measured at the sample mount during cooldown of the setup. Vertical dashed line marks the moment of the end of film boiling of the copper rod, visible as sharp negative spike in the temperature curve after ~2,700 s. Temperature stabilises after ~5,500 seconds at -189 °C. Afterwards the setup start warming up at a rate of 12 mK/min. Measurements can be performed for ~12 h before having to refill the $LN_2$ dewar of the setup.

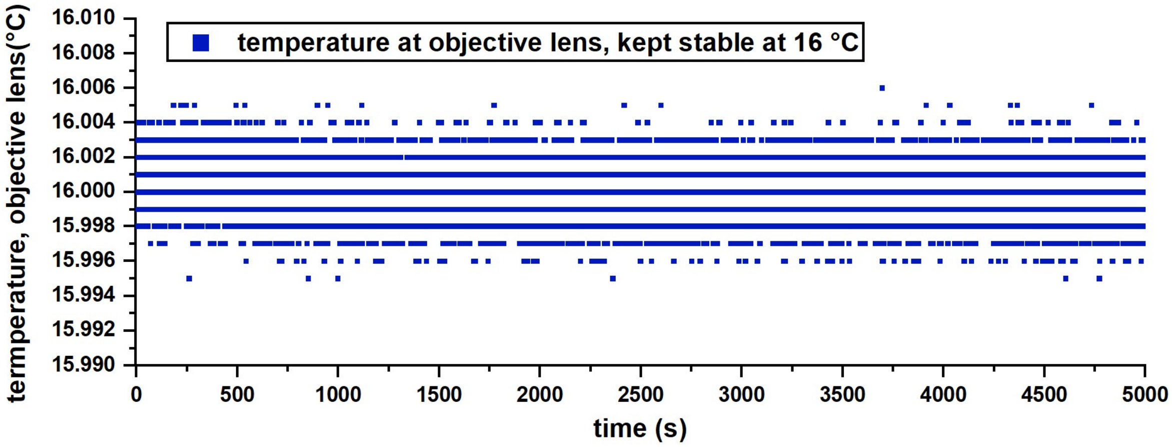


**Figure S2 | Temperature at objective lens during measurements.** Temperature of objective lens was set to 16.0 °C. The temperature was recorded with a calibrated Cernox resistive sensor (LS-CX-1080-SD-HT-20M_P, LakeShore). The difference between the set and the average measured temperature is only 0.2 mK (with standard deviation of 2.3 mK over the course of 5000 s). To compensate for the slow heating of the setup after cooldown (see suppl. Fig S1), the temperature of the objective lens has to be increased by 2 mK/min.

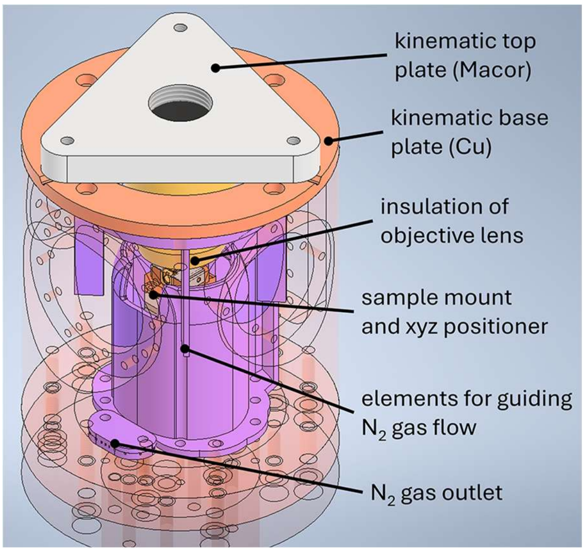


**Figure S3 | VUCROM cryo-chamber.** The heated and insulated objective lens is mounted in a Macor plate that form the top of a kinematic mount. Set screws with ball tips allow angle adjustment of the objective lens. The Macor plate thermally decouples the objective lens from the kinetic base plate and the housing of the cryo chamber, which are made from copper. Additional elements (purple) in the cryo-chamber built around the sample mount, xyz-positioner and the objective lens enhance a laminar flow of the $N_2$ gas to minimise vibrations.

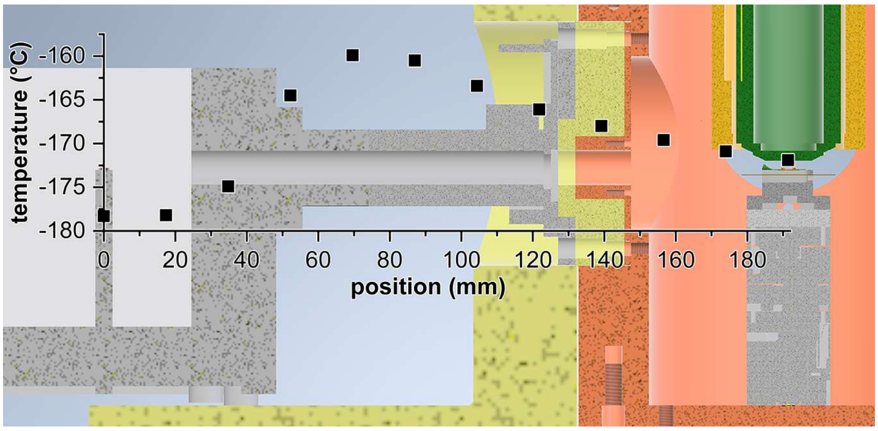


**Figure S4 | Temperature of sample cartridge during transfer.** Schematic illustration of Leica Transfer Shuttle, docking port at setup and inside of cryo-chamber with sample positioner and objective lens (from left to right). Position 0 mm corresponds to loading position of cartridge in the Transfer Shuttle. At position 192 mm the cartridge has been placed into the sample holder on the xyz-positioner. Temperature was measured with a thermocouple mounted to the gripper of the Leica Transfer Shuttle.

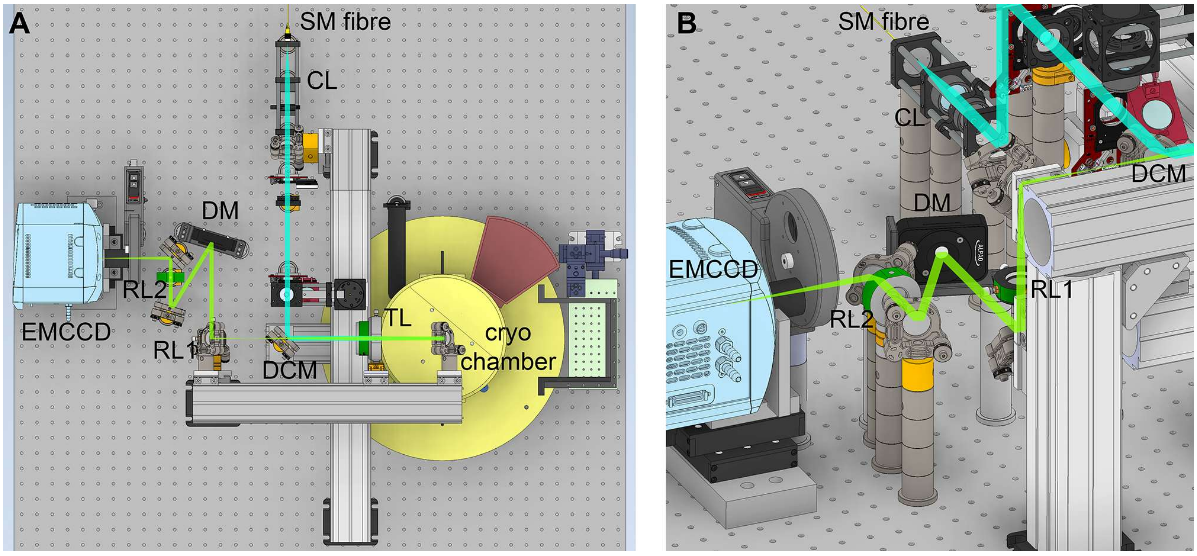


**Figure S5 | Integration of deformable mirror (DM) in cryo super-resolution FM setup. A**: Schematic top view of microscopy system with excitation (cyan) and detection (green) paths indicated. SM fibre: single mode fibre, CL: collimation lens, TL: tube lens, DCM: dichroic mirror, RL1: relay lens 1 (160 mm), RL2: relay lens 2 (160 mm). DM placed at the conjugate plane to the back focal plane of the objective lens. Incident angle of the detection light to the plane of the DM: ~17°. Objective lens (0.9 NA, Nikon MUC11900) and sample stage are located inside the cryo chamber (for details see Fig. 1). **B**: Side view of detection path with DM.

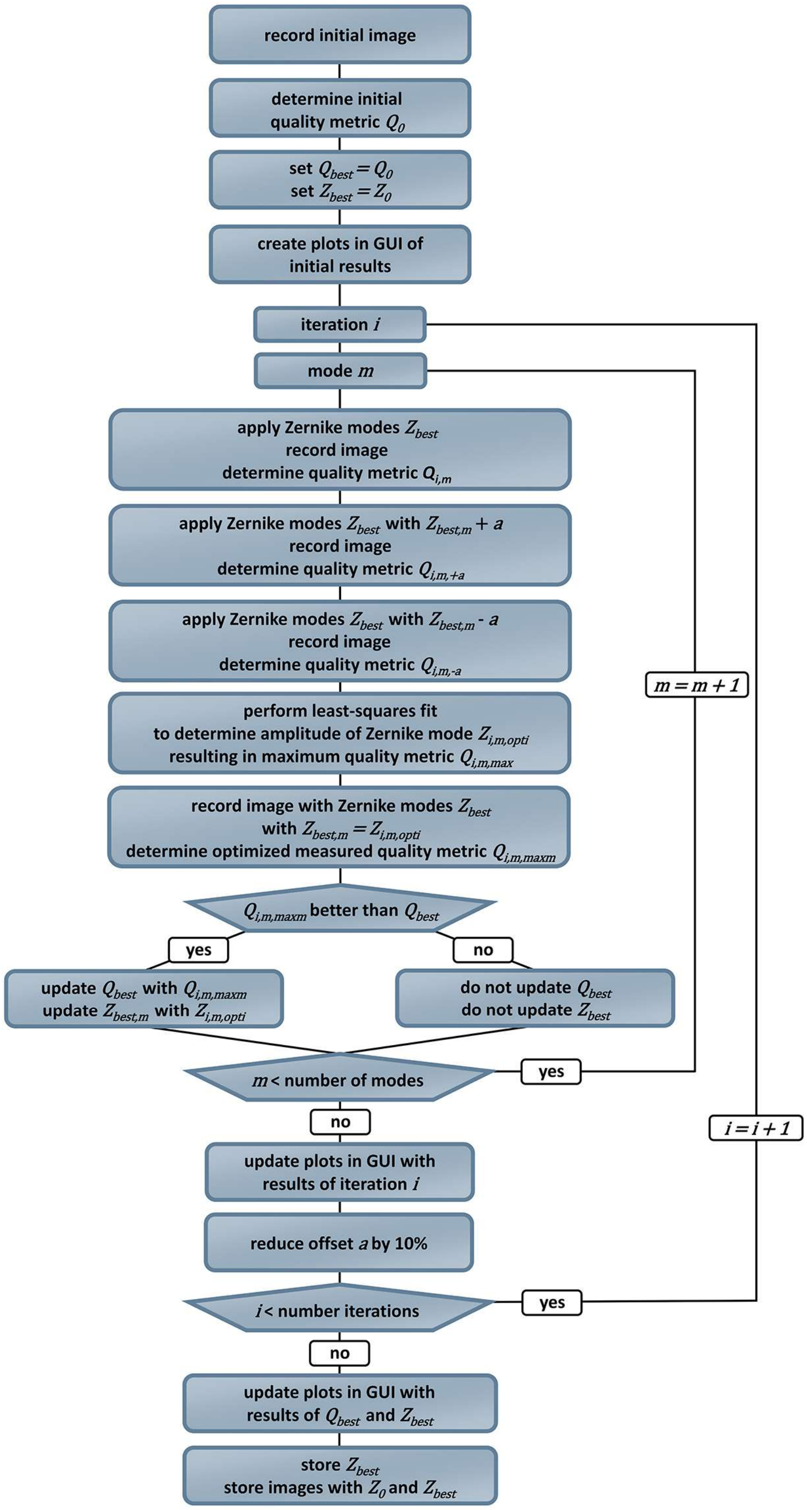


**Figure S6 | Schematic overview of sensorless AO correction routine developed for cryo-FM.** The correction routine follows the general framework for sensorless AO correction as described in McFadden *et al.* [1]. Our AO correction routine for cryo-FM first determines the initial quality metric $Q_0$ for Zernike modes $Z_0$ applied to the DM that correspond to a flat mirror. The quality metric $Q$ is defined as:

$$Q = \frac{I_{cent}}{c_I} \frac{1}{c_\omega \ \omega} \frac{1}{c_A + \sqrt{A_x^2 + A_y^2}}$$

with central intensity $I_{cent}$ as measured in the central 3x3 pixels of point spread function (PSF), width of PSF $\omega$ as measured at 20% of maximum intensity and asymmetries of the PSF $A_x$ and $A_y$ in x- and y-direction. Asymmetry is measured by comparing the centroids determined when either restricting the PSF fitting procedure (adapted from Grüll *et al.* [2]) to a smaller area or allowing a larger area around the PSFs (radius of 700 nm vs. 1,300 nm).

The latter is more sensitive to the periphery of the PSF and is more affected by aberrations distorting the PSF asymmetrically, e.g. coma. The contribution of each quality metric can be adjusted by the corresponding weighting factors $c_I$, $c_\omega$ and $c_A$. In each iteration $i$, quality metric $Q_{i,m}$ is determined for every Zernike mode $m$ as stored in $Z_{best}$ corresponding to the best measured quality metric $Q_{best}$ and additionally with Zernike modes with positive and negative offsets $a$. Based on the measured pairs $Z_{best}$ and $Q_{i,m}$; $Z_{best,m,+a}$ and $Q_{i,m,+a}$; $Z_{best,m,-a}$ and $Q_{i,m,-a}$, a least-squares fit determines the optimised amplitude for Zernike mode $m$: $Z_{i,m,opti}$, for which the corresponding quality metric has its maximum value $Q_{i,m,max}$. A new image is recorded with Zernike modes applied to the DM corresponding to $Z_{best}$ with $Z_{best,m} = Z_{i,m,opti}$. The corresponding measured quality metric $Q_{i,m,maxm}$ is determined. $Q_{i,m,maxm}$ is stored as the new $Q_{best}$ if its value is better than $Q_{best}$. In this case the mode $m$ in $Z_{best}$ is updated by $Z_{i,m,opti}$. This procedure is repeated for every Zernike mode $m$ and iterated $i$ times as defined by the user. Zernike modes can be limited by the user to only primary modes to speed up the procedure if only a rough estimate is required.

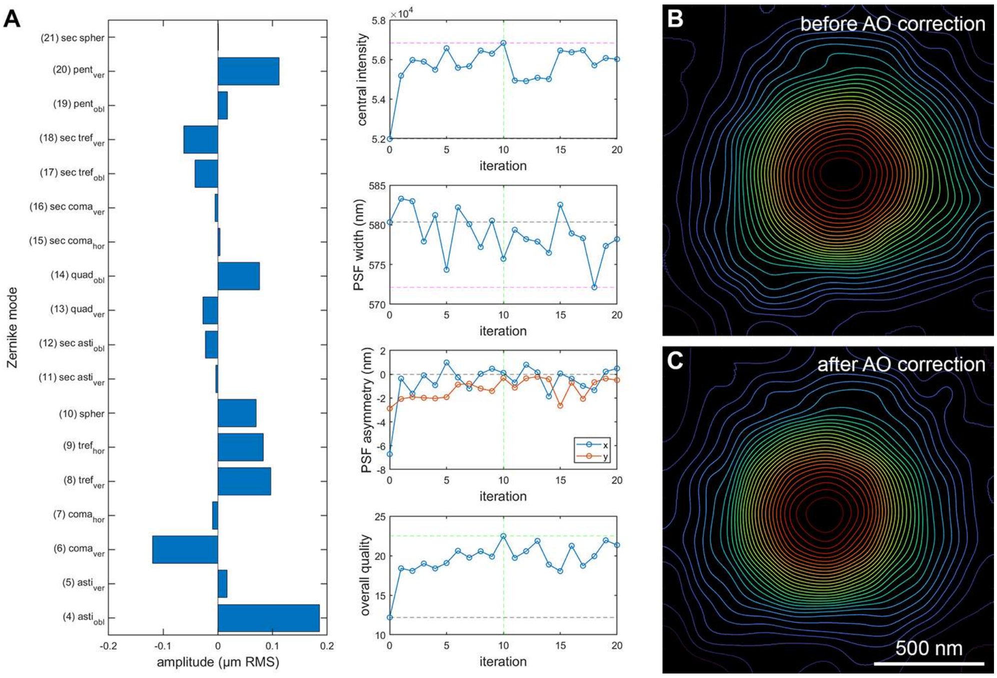


**Figure S7 | AO correction of aberrations inherent to the super-resolution cryo-FM setup. A**: Left diagram displays amplitudes of Zernike modes resulting in highest score of the overall quality metric after 20 iterations of AO correction routine for optically isolated fluorescent beads. Plots on the right show the results of the different individual quality metrics: intensity, PSF width, PSF asymmetry and the overall metric. The parameters resulting in the highest overall score (green dashed lines) are kept as the best approximation for minimizing the present aberrations. The individual quality metrics show that measured intensity, which is a common metric for AO routines, only changes by ~10% between before and after AO correction. In comparison, the metric assessing asymmetries in the PSF changes by ~10-fold, even for the low level of aberrations in this case. **B, C**: Contour plots of PSF of one bead before (**B**) and after (**C**) AO correction based on the parameters shown in **A**. Intensity is displayed on a logarithmic scale to increase visibility of differences at the periphery of the PSF.

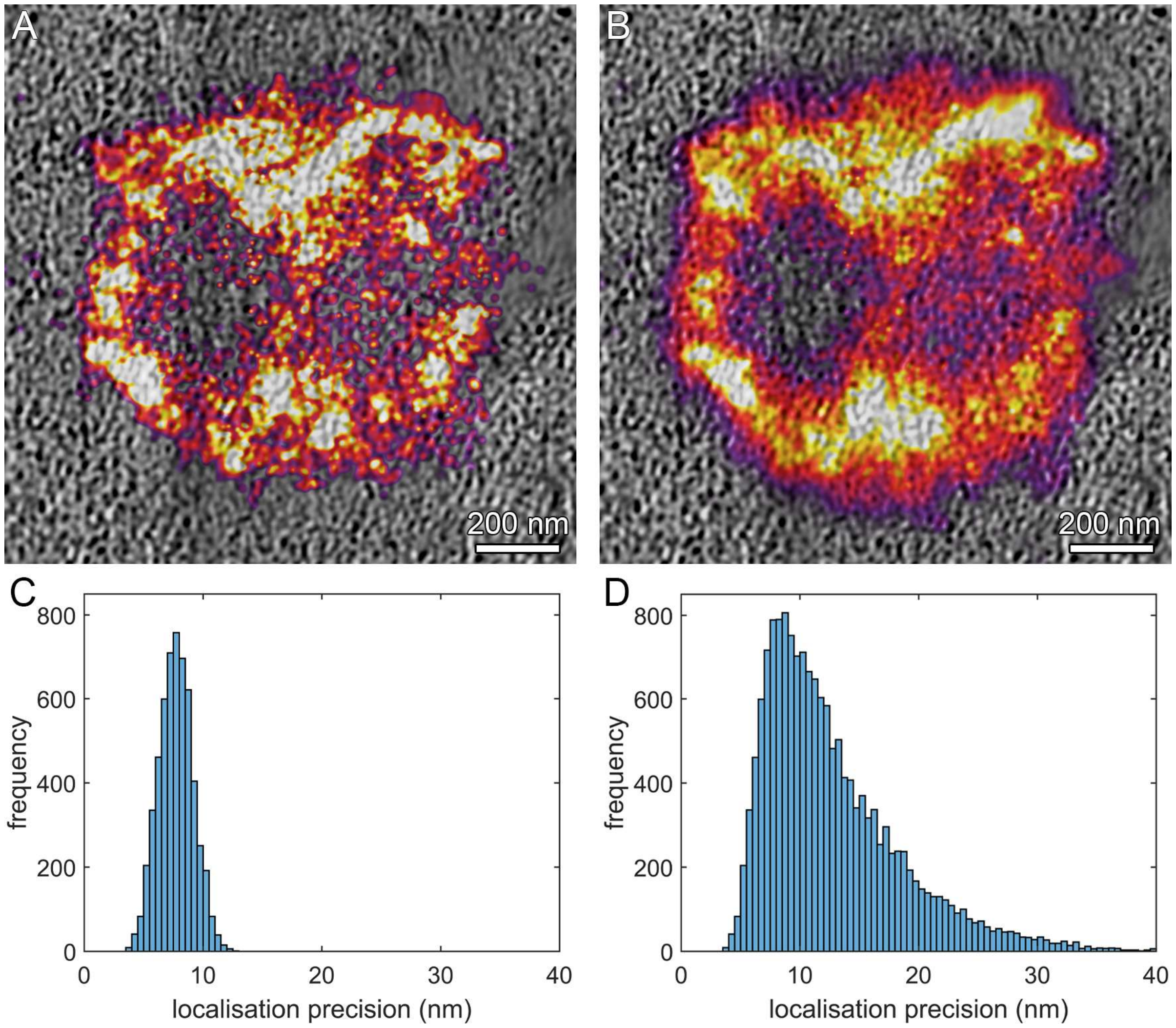


**Figure S8 | Super-resolution cryo-CLEM of YFP-labelled PML body with different thresholds for single molecule detection. A**: Cryo-SMLM reconstruction of same PML body as in image shown in Fig. 5D (cryo-SMLM overlay with projection of overview tomogram, nominal magnification: 4,800×). Threshold for single molecule localisation was $18\ \sigma_B$ with $\sigma_B = \sqrt{N_B}$ and $N_B$ being the background intensity. **B**: Cryo-SMLM reconstruction of the same PML body as shown in **A**, but with a threshold for single molecule localisation of $4\ \sigma_B$. **C**: Histogram of localisation precisions for the single molecules detected as shown in **A**. Average single molecule localisation precision: 7.7 nm. Total amount of detected signals: 5,509. **D**: Histogram of localisation precisions for the single molecules detected as shown in **B**. Average single molecule localisation precision: 12.7 nm. Total amount of detected signals: 16,113.

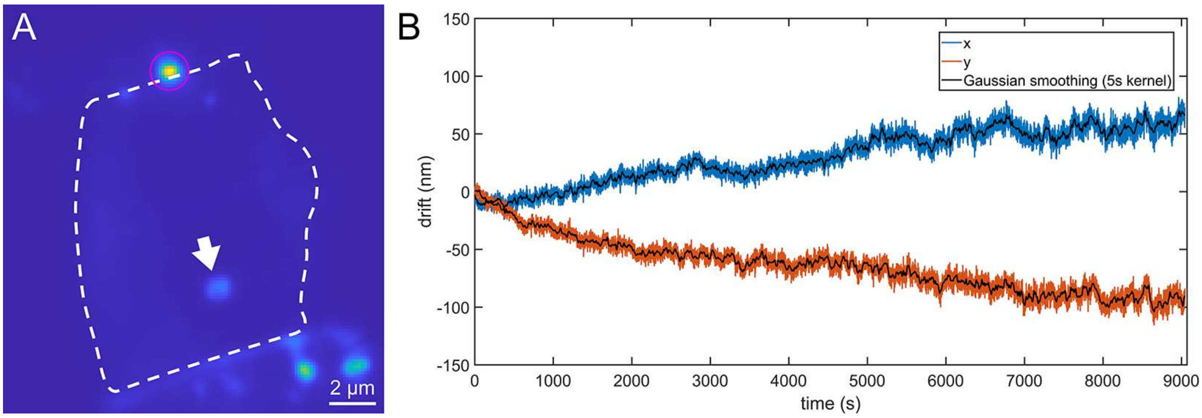


**Figure S9 | Drift correction for cryo-SR-FM imaging of YFP-labelled PML body. A**: Bright object (marked by purple ring) with no observable photo-blinking outside of lamella (dashed lines indicate outline of lamella) was selected to determine and correct drift during cryo-SR-FM measurement. Arrow indicates PML body shown in cryo-SR-CLEM images in Fig. 4 and Fig. S8. **B**: Drift of the object marked in **A** over the course of the entire cryo-SR-FM measurement with 500 ms time steps. Determined drift was smoothed with a 5 s wide Gaussian for drift correction of the cryo-SR-FM data to minimise variations due to localisation precision limited by number of available photons.

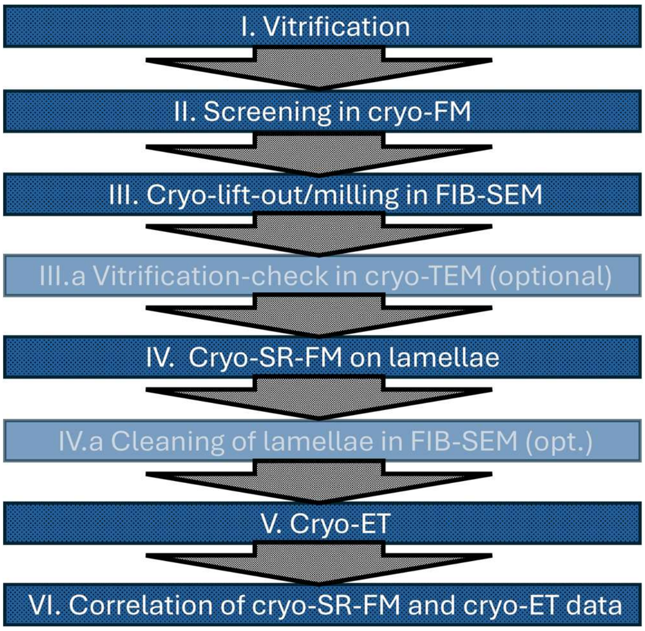


**Figure S10 | On-lamella cryo-SR-CLEM workflow. I**: Vitrification of specimen using plunge-freezing or high pressure freezing. **II**: Screening of vitrified specimens by cryo-fluorescence microscopy for selecting FIB-milling positions. For thick specimens (e.g. high-pressure-frozen tissue) a confocal system is recommended as it offers optical sectioning; for sparse weakly fluorescent/labelled structures a wide-field setup is recommended as it offers higher sensitivity. **III**: Cryo-lift-out/FIB-milling according to information provided by cryo-fluorescence imaging (**II**). Milling of additional markers such as crosses (Fig. 5) allows achieving higher correlation precision. **III.a**: Vitrification of lamellae should be checked before starting cryo-SR-FM measurements by TEM (low magnification, e.g. 2,000-4,000× and low dose ~1 $e^{-}Å^{-1}$). **IV**: Cryo-SR-FM data acquisition. **IV.a**: Optional cleaning of lamellae from ice contaminations gathered during the repeated transfer steps. Ice contamination can be reduced by using a glove box with close to zero humidity for transfer/handling steps. **V**: Cryo-ET data acquisition in lamellae of features imaged by cryo-SR-FM (**IV**) and cryo-ET processing. **VI**: Correlation of cryo-SR-FM and cryo-ET data.
